# Dietary preservatives alter the gut microbiota *in vitro* and *in vivo* with sex-specific consequences for host metabolic development

**DOI:** 10.1101/2024.05.14.593600

**Authors:** Laura D. Schell, Katia S. Chadaideh, Cary R. Allen-Blevins, Emily M. Venable, Rachel N. Carmody

**Author notes:** Correspondence: Laura D. Schell, Department of Human Evolutionary Biology Harvard University, 11 Divinity Avenue, Cambridge, MA 02138, Rachel N. Carmody, Department of Human Evolutionary Biology Harvard University, 11 Divinity Avenue, Cambridge, MA 02138. **Classification**: Biological Sciences / Physiology.

## Abstract

Antibiotics in early life can promote adiposity via interactions with the gut microbiota. However, antibiotics represent only one possible route of antimicrobial exposure. Dietary preservatives exhibit antimicrobial activity, contain chemical structures accessible to microbial enzymes, and alter environmental conditions favoring specific microbial taxa. Therefore, preservatives that retain bioactivity in the gut might likewise alter the gut microbiota and host metabolism. Here we conduct *in vitro, ex vivo, and in vivo* experiments in mice to test the effects of preservatives on the gut microbiota and host physiology. We screened common dietary preservatives against a panel of human gut isolates and whole fecal communities, finding that preservatives strongly altered microbial growth and community structure. We exposed mice to diet-relevant doses of 4 preservatives [acetic acid, BHA (butylated hydroxyanisole), EDTA (ethylenediaminetetraacetic acid) and sodium sulfite], which each induced compound-specific changes in gut microbiota composition. Finally, we compared the long-term effects of early-life EDTA and low-dose antibiotic (ampicillin) exposure. EDTA exposure modestly reduced nutrient absorption and cecal acetate in both sexes, resulting in lower adiposity in females despite greater food intake. Females exposed to ampicillin also exhibited lower adiposity, along with larger brains and smaller livers. By contrast, in males, ampicillin exposure generally increased energy harvest and decreased energy expenditure, resulting in higher adiposity. Our results highlight the potential for everyday doses of common dietary preservatives to affect the gut microbiota and impact metabolism differently in males and females. Thus, despite their generally-regarded-as-safe designation, preservatives could have unintended consequences for consumer health.

**SIGNIFICANCE:** Early-life exposure to antibiotics can alter the gut microbiota and shape adult metabolic health. Here we show that dietary preservatives can have analogous effects. Common dietary preservatives altered gut microbiota composition *in vitro*, *ex vivo* and *in vivo*. Early-life EDTA exposure had long-term, sex-specific consequences for energy metabolism. Simultaneously, we deliver new mechanistic understanding of early-life antimicrobial-induced effects on adiposity via evidence that low-dose ampicillin treatment increases energy harvest while conserving energy expenditure in males, promoting adiposity, while EDTA treatment dampens energy harvest, promoting leanness in females. Overall, our results emphasize that early-life gut microbiome disruptions can be triggered by diverse antimicrobial exposures, with previously unappreciated metabolic consequences that differ for males and females.

## INTRODUCTION

Humans have long practiced diverse food preservation methods to extend shelf life and prevent food spoilage. Drying, salting, smoking, and fermentation all function by making food an inhospitable environment to undesirable microbes, either by removing water or adding compounds with antimicrobial activity against foodborne microbes. The ubiquity of traditional or industrial preservatives in human diets, combined with their activity against foodborne bacteria raise the possibility that, once consumed, preservatives might also affect some of the trillions of microbes living in the gastrointestinal tract.

Gastrointestinal microbes (collectively, gut microbiota) are far from passive inhabitants. The gut microbiota plays critical roles in nutrient digestion, energy allocation, immunological training and maintenance, and neurologic and endocrine activity^1–4^. Consequently, variation in the gut microbiota can affect growth, development, and many gastrointestinal, autoimmune, neurological, and metabolic diseases^5^. Both diet and antibiotic use can rapidly re-shape the gut microbiota, whether by encouraging growth of some taxa at the expense of others or else by altering the functions carried out by different microbes within the gut.

Differences in dietary macronutrient content^6^, plant versus animal sources^7^, cooking^8^, and fermentation^9^ can all reshape the gut microbiota, often with downstream effects on host metabolism. Non-nutritive dietary compounds – such as phytochemicals and food additives – can also affect the gut microbiota with consequences for host health^10–12^. For instance, consumption of dietary emulsifiers by mice at diet-relevant doses altered the gut microbiota and promoted obesity and insulin resistance^12^. Importantly, germ-free mice were protected from these effects, and effects were transmissible to germ-free mice upon gut microbiota transplantation, suggesting that emulsifier-induced changes in the gut microbiota were necessary and sufficient to link emulsifier ingestion to biomarkers for metabolic syndrome^12^. Many drugs can also influence the gut microbiota^13,14^, with antibiotics being a canonical example. When administered to mice even at very low, subtherapeutic levels, antibiotics have been shown to alter gut microbiota composition and, when treatment starts early in life, to promote weight gain and increased body fat in adulthood^15–17^.

Given the antimicrobial activity of preservatives and the potential for even low doses of antimicrobials to alter the gut microbiota, preservatives represent a key, largely unexplored influence on health. In this study, we used *in vitro* and *in vivo* approaches to test the effect of common dietary preservatives on the gut microbiota and to assess potential consequences for host metabolism. We first screened 9 common dietary preservatives [acetic acid, BHA, BHT (butylated hydroxytoluene), EDTA, sodium benzoate, sodium chloride, sodium nitrate, sodium sulfite, sulfur dioxide] for activity against a small panel of human gut isolates (*Bacteroides ovatus*, *Clostridium symbiosum*, *Eggerthella lenta*, *Escherichia coli*), representing 4 of the most abundant phyla in the human gut. We then tested the effects of a subset of these compounds (acetic acid, BHA, EDTA, sodium sulfite) on whole gut microbial communities both *ex vivo* and in mice. Last, we investigated the effects of long-term and early-life exposure of one preservative, EDTA, on the developing microbiota and long-term host metabolism. Most of the preservatives tested exhibited strong activity against gut bacteria *in vitro*, *ex vivo*, and in mice. Exposure to EDTA starting in early life had pronounced effects on the developing gut microbiota and increased fecal energy density. For females, EDTA exposure resulted in altered energy balance, as evidenced by reduced fat mass and feed efficiency in treated females versus untreated female controls. Overall, our results indicate that dietary preservatives can alter the gut microbiota and that long-term exposure to preservatives starting early in life, as occurs in many human populations, may carry metabolic consequences.

## RESULTS

### Dietary preservatives alter the growth of individual gut isolates and whole communities in vitro

We first evaluated the antimicrobial activity of 9 common dietary preservatives at diet-relevant concentrations against a panel gut bacterial isolates under anaerobic conditions (Table S1). Almost all compounds, with the exception of sodium chloride and sodium nitrate, significantly inhibited bacterial growth at or below the concentrations used in food (Figure S1). BHA and EDTA were notable for their antimicrobial activity against all strains at and below the maximum concentrations allowed in food (200 µg/ml and 1000 µg/ml, respectively). Of the 2 traditional preservatives, acetic acid (vinegar) and sodium chloride (table salt), acetic acid was of particular interest because it had divergent effects at higher concentrations, promoting growth of *C. symbiosum* while inhibiting growth of *E. coli* and *B. ovatus*. The 2 sulfur-based compounds tested, sodium sulfite and sulfur dioxide, showed similar patterns of growth inhibition starting at the maximum concentrations allowed in food (500 µg/ml).

To elucidate the broader antimicrobial effects of preservatives exhibiting substantial inhibitory effects on single isolates, we next tested 4 preservatives (acetic acid, BHA, EDTA, sodium sulfite) on whole gut bacterial communities cultured *ex vivo*, with the broad-spectrum antibiotic ampicillin as a positive control. Whole mouse fecal communities (n=4 donor mice) were grown in differing concentrations of each compound for 48 hours, with monitoring of overall growth by optical density (OD_600_), confirmation of cell density measurements by plate culture (Figure S2), and profiling of community composition at endpoint by 16S rDNA sequencing. As expected, ampicillin altered microbial growth dynamics and endpoint community composition at all concentrations tested. All 4 preservatives also significantly inhibited maximum community growth (Figure 1A) and altered community composition (Figure 1B-C), with clear dose-response effects. These effects were strongest for BHA and EDTA, both of which significantly altered community composition at the lowest concentration sequenced (50 µg/ml), well below the maximum concentrations allowed in food (Table S1). Where there was undetectable cell growth (as measured by OD_600_, validated with CFU counts, Figure S2), community composition resembled the original inocula likely because that was the only DNA present in the sample (Figure 1).

**Figure 1.**
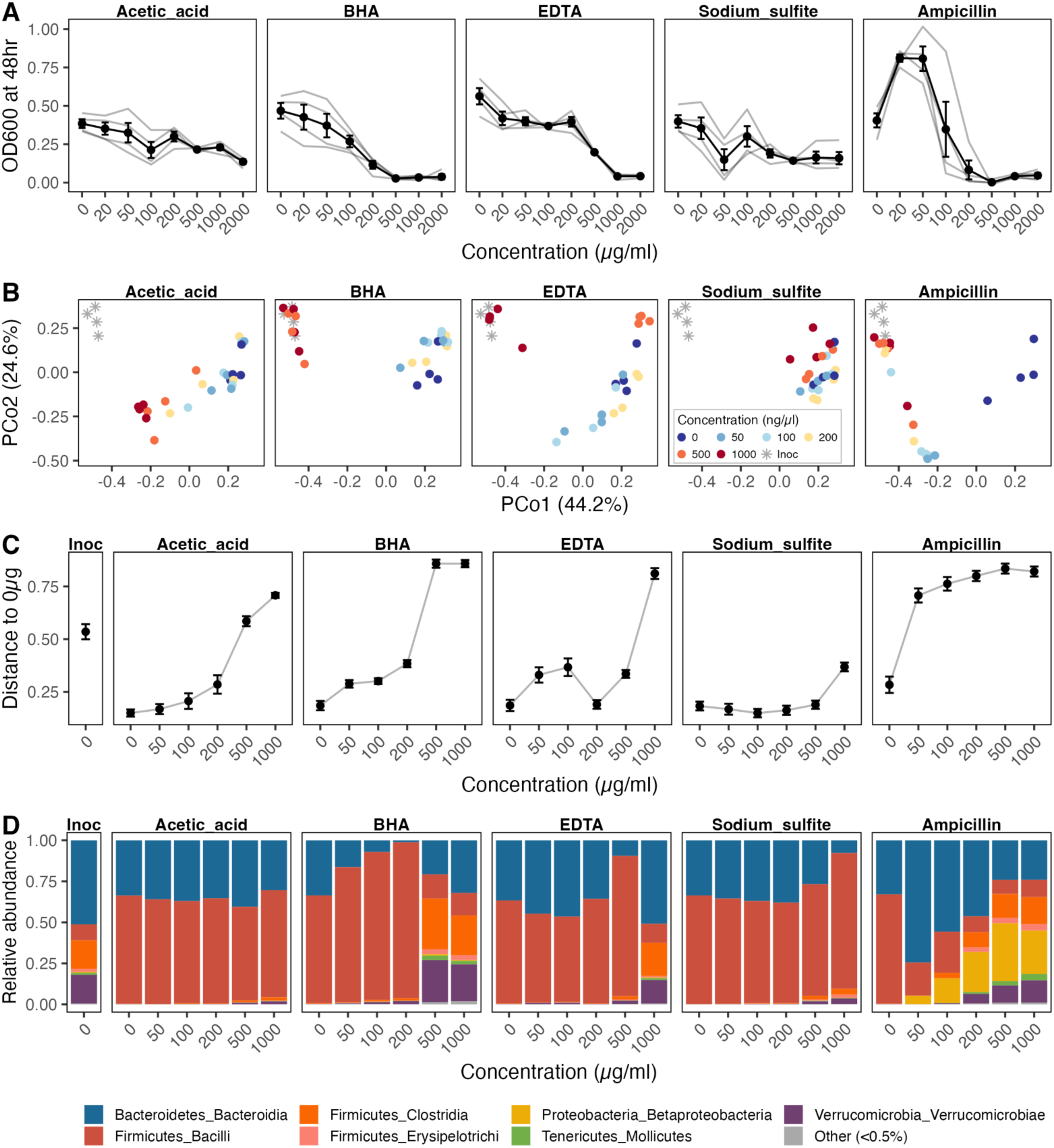
Impact of preservatives on whole gut communities *ex vivo* after 48 hours of growth. (A) Cell density at endpoint of whole mouse gut microbial communities grown in media containing varying concentrations of each preservative. Data are mean ± SE, with gray lines indicating each of 4 biological replicates. (B) Principal coordinate plot representing dissimilarity among *ex vivo* microbial communities, as indexed by Bray-Curtis distance. The original inoculum is indicated as a gray star. (C) Bray-Curtis distance between each community and the associated no-compound control. Larger values indicate a greater impact of the preservative on microbial community composition. Data are mean ± SE. (D) Mean taxonomic composition of each microbial community, showing relative abundance by class.

At concentrations where OD_600_ readings indicated uninhibited or minimally inhibited growth, each preservative altered community composition in slightly different ways. Low doses of ampicillin greatly favored several prevalent strains of Bacteroidetes and Proteobacteria, to the extent a higher maximum OD_600_ was observed at 20 and 50 µg/ml. Among the preservatives, increasing concentrations of BHA reduced the relative abundance of Bacteroidetes in favor of Firmicutes, with a similar trend trend for higher levels of EDTA. Principal coordinate analysis also highlighted the impact of increasing concentrations of each preservative (Figure 1B), particularly for acetic acid, where the statistically significant effects of dose are readily observable in PCoA space but harder to visualize in terms of relative abundance.

### Short-term preservative exposure has compound-specific effects on the gut microbiota of adult mice

To examine how dietary preservatives affect the gut microbiota *in vivo*, we treated adult male C57BL6/J mice with one of each preservative for 7 days via drinking water, with doses set at the maximum acceptable daily intake listed by the FDA (see Table S1) and scaled isometrically for mice (see Methods). An additional group treated with a low dose of ampicillin (6.7 µg/ml) was included as a positive control, along with an untreated water group. Water intake was measured every 2-3 days to confirm dosing estimates. Neither ampicillin nor any preservative significantly affected mouse body mass during this short-term exposure, with the exception of the acetic acid, where mouse water intake dropped by about 1/3 during treatment, likely due to an aversion to the vinegar taste (Figure S3).

We used 16S rDNA sequencing to profile the fecal microbiota daily, as well as the gut microbiota along the length of the gastrointestinal (GI) tract at endpoint. The strongest overall determinant of gut microbiota composition, as indexed by Bray-Curtis distance, in both fecal and gut effluent samples was the original cage in which mice were housed prior to redistribution into individual housing at the start of the study (p<0.001, R^2^=0.311–0.549, PERMANOVA) (Figure S4, Table S2). Microbial communities at endpoint also differed significantly by GI tract location (p<0.001, R^2^=0.272). Therefore, in all microbiome analyses, we controlled for the effects of cage and sample location, as the PERMANOVA test ascribes variation sequentially and failing to account for these high-impact variables can prevent the detection of biologically significant differences by treatment.

As expected, low-dose ampicillin treatment altered gut microbiome composition, both longitudinally in fecal samples (p=0.002, R^2^=0.033, PERMANOVA, Table S2) and at endpoint along the GI tract (p=0.003, R^2^=0.034, PERMANOVA) without any detectable impact on bacterial density (Figure S5). All 4 tested preservatives also influenced gut microbial community composition over time in at least 2 of 3 different longitudinal models (Table S2). This effect was particularly robust for EDTA, where the longitudinal effect of treatment remained significant across all 3 models tested, including after the addition of a ‘Phase’ variable to control for stochastic variation between baseline and treatment periods that may be shared with control mice (EDTA treatment: p<0.05, R^2^=0.016–0.026). In contrast, preservatives had no effect on microbiota composition in any cross-sectional analyses of endpoint samples, even when including samples from all GI tract locations, potentially due to high variability by GI location and inability of cross-sectional analyses to use within-mouse controls.

We next used MaAsLin2^18^, a tool that applies general linear models to determine multivariate associations with metagenomic features, to identify microbial taxa that were particularly susceptible or resistant to treatment with dietary preservatives. Again, we were able to compare the effects of treatment longitudinally (with current treatment status as a fixed effect and source cage as a random effect) and cross-sectionally (with treatment and sample location as fixed effects and source cage as a random effect). These models identified a large number of differentially abundant taxa at multiple taxonomic levels that were impacted by preservative use (Figure 2). Among preservatives, there was some overlap regarding the microbial taxa that were consistently altered, such as the genus *Clostridium* (family: Ruminococcaceae), which was reduced in all treatment groups in either endpoint or longitudinal models. In many other cases, preservatives had compound-specific effects: for instance, the relative abundance of *Allobaculum*, a genus recently implicated in the attenuation of insulin resistance^19^, was elevated by EDTA treatment but reduced by acetic acid, sodium sulfite, and ampicillin. Overall, each preservative treatment had a unique impact on the gut microbiome, characterized by subtle shifts in overall composition and distinct differentially abundant taxa.

**Figure 2.**
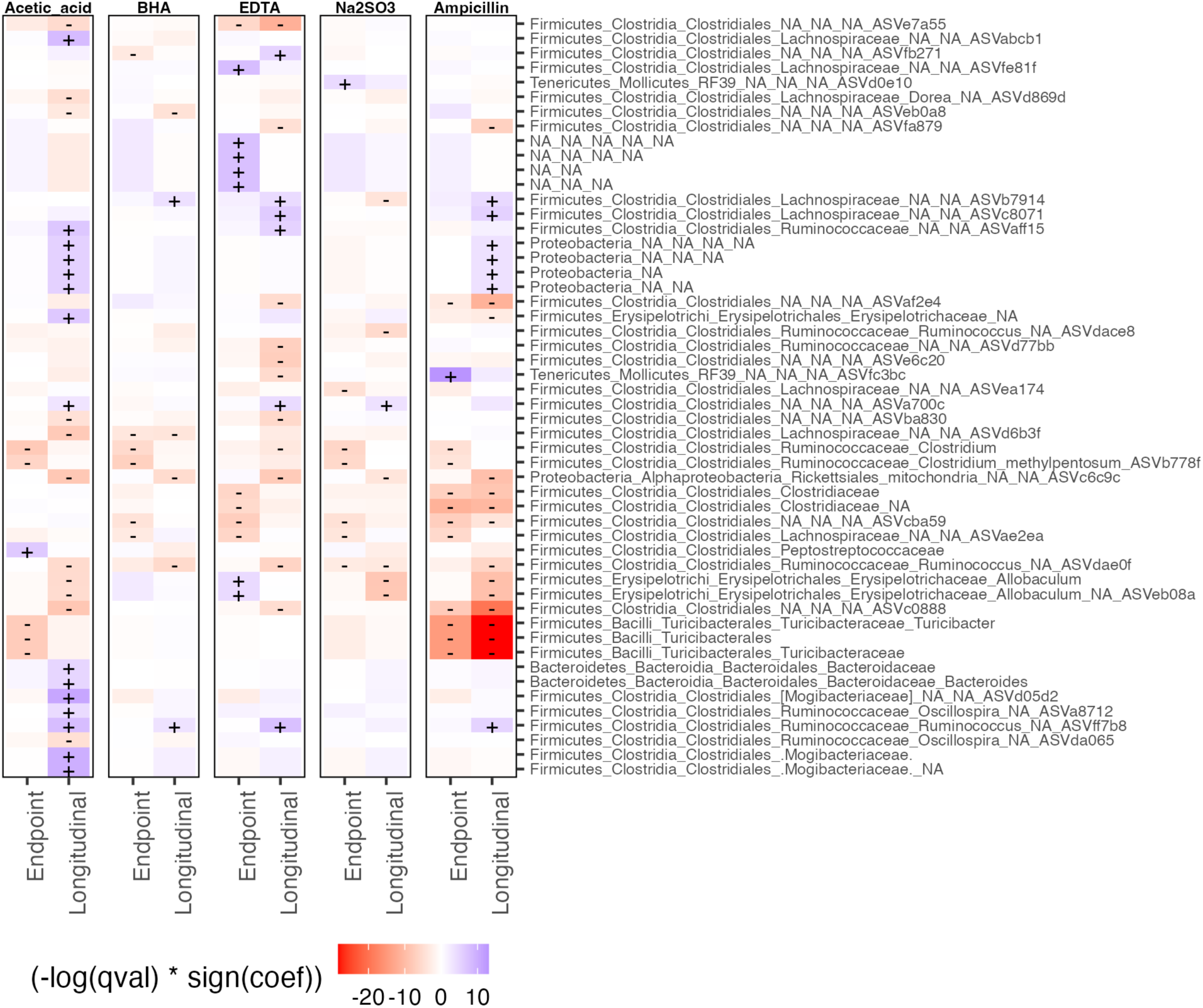
Differentially abundant taxa in 7-day *in vivo* trial identified using MaAslin2. Results displayed from 2 different models, endpoint and longitudinal, showing results from all taxonomic levels (phylum through ASV) with corresponding taxonomic classification, where available. The endpoint model was run on data from all endpoint (Day 7) samples, including the 4 points sampled along the GI tract, with treatment and GI location as fixed effects and source cage as a random effect. The longitudinal model captured fecal samples from baseline (Day –2) through treatment Day 6 and used treatment as a fixed effect with source cage as a random effect. Direction and strength of effect are indicated by color, with statistically significant effects (q<0.05) indicated by + or – signs. NA indicates taxonomic levels with no known classification.

### Dietary antimicrobials alter murine gut microbiota strongly in early life

The gut microbiota changes over the course of host development, and perturbations of the gut microbiota during gestation and infancy—including with low-dose antibiotics—have previously been shown to have long-term effects on host metabolism^16^. To examine how preservatives with antimicrobial properties might specifically affect the developing gut microbiota and host metabolism, we treated pregnant mice and their litters with either diet-relevant doses of EDTA, low-dose ampicillin (positive control), or normal drinking water (negative control) from gestational day 13.5 until offspring were 28 weeks old.

Consistent with previous studies^15,16^, low-dose ampicillin treatment altered gut microbiota composition for the duration of treatment (Figure 3A, Table S3), with reductions in a number of individual microbial taxa identified using MaAslin2 (Figure 3B). The genus *Allobaculum* was again notable among these differentially abundant taxa (Figure S6) as it is nearly absent in ampicillin-treated mice, was present at high levels in untreated mice during early life, and has also shown consistent reductions by low-dose antibiotic treatment in prior work^16^. For other genera such as *Clostridium* (family: Ruminococcaceae), EDTA but not ampicillin treatment reduced abundance, particularly during early life (Figure S6), again indicating that the preservatives are not simply less potent antimicrobials than antibiotics, but may also have distinct impacts on gut microbes.

**Figure 3.**
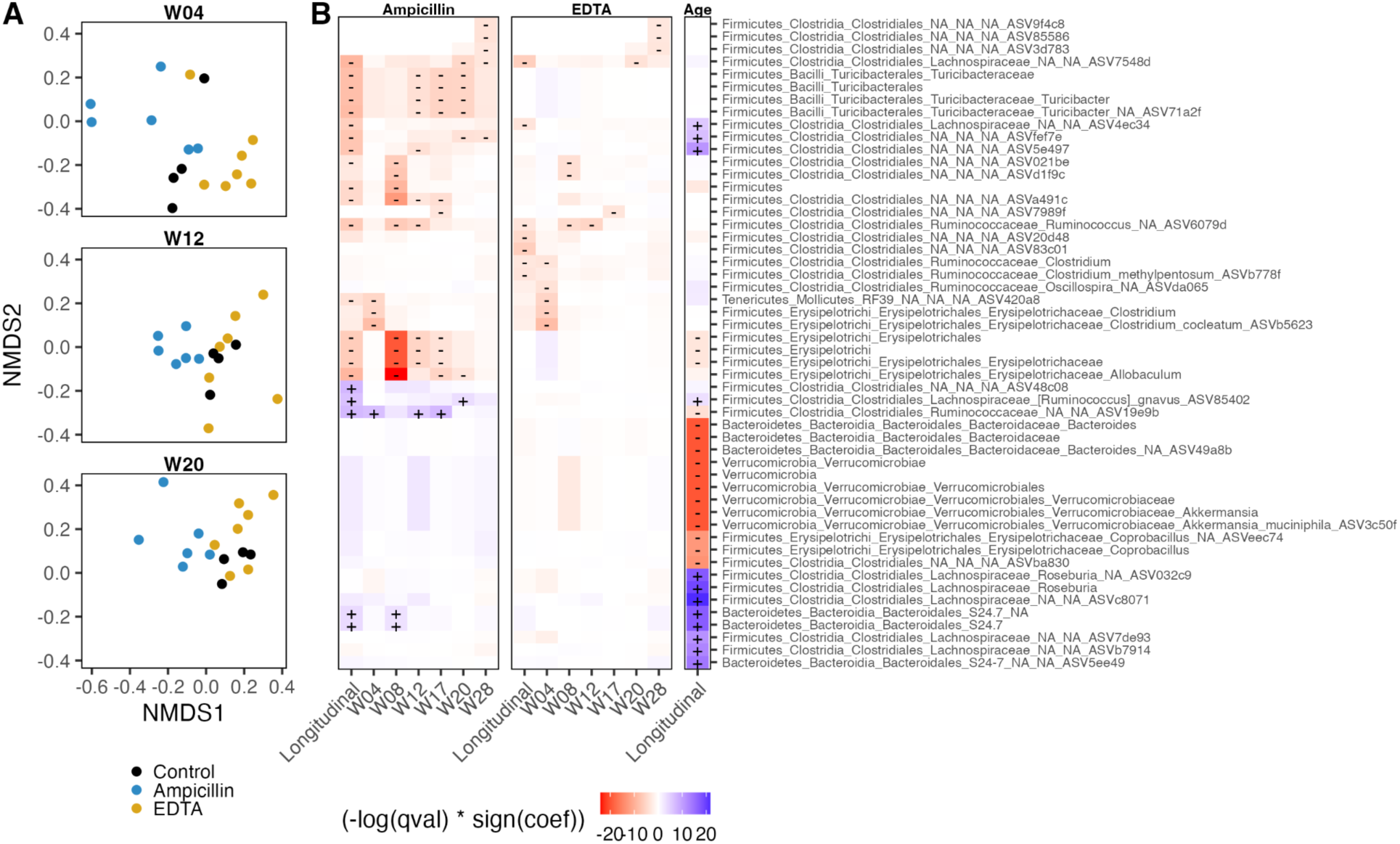
Gut microbiota composition in long-term trial. (A) Non-dimensional linear scaling (NMDS) ordination plot of Bray-Curtis distances between mouse gut microbiomes at 4, 12, and 20 weeks of age. Top 50 taxa that differ significantly by treatment and/or age, as identified by MaAslin2. Models were run across all ages with ‘Age’ as an additional fixed effect and ‘Mouse ID’ as a random effect (longitudinal model) or else at each age independently (W04 through W28). Direction of effect is indicated by color, with darker shades indicating lower q values and statistically significant effects (q<0.05) indicated by + or – signs.

Treatment with EDTA also significantly altered gut microbiota composition, but this effect was largely limited to early life, with a significant effect of EDTA treatment at 4 weeks of age (p=0.037, PERMANOVA, Table S3) but not for timepoints between 8 and 20 weeks of age (p=0.552–0.885). The amount of overall variation ascribed to EDTA treatment also generally declined from week 4 (R^2^=0.179) into later weeks (R^2^=0.058–0.080). Consequently, while MaAsLin2 identified 17 taxa as differing significantly with EDTA treatment (Figure 3B), most of these were either reduced only at 4 weeks or else only rose to significance in the longitudinal model that included mouse age as an additional variable. Overall, our data indicate that young mice may be particularly sensitive to EDTA-induced perturbations of the gut microbiota or else that the gut microbiota can become resistant to the effect of EDTA with prolonged treatment.

### Early-life antimicrobial treatment alters host adiposity and energy gain

Because of the role of the gut microbiome in programming host metabolism^20^, we tracked body composition and energy intake over the course of the experiment. Notably, early-life exposure to either low-dose ampicillin or EDTA had sex-specific outcomes. Ampicillin treatment starting in gestation resulted in increased body fat in males, whether measured as total body fat via EchoMRI (Figure 4A) or the mass of the epididymal fat pad at 28 weeks (Figure 4B). Intriguingly, however, both EDTA and ampicillin-treated females exhibited decreased body mass as adults compared to untreated controls, with reductions coming largely from body fat rather than lean mass (Figure 4A).

**Figure 4.**
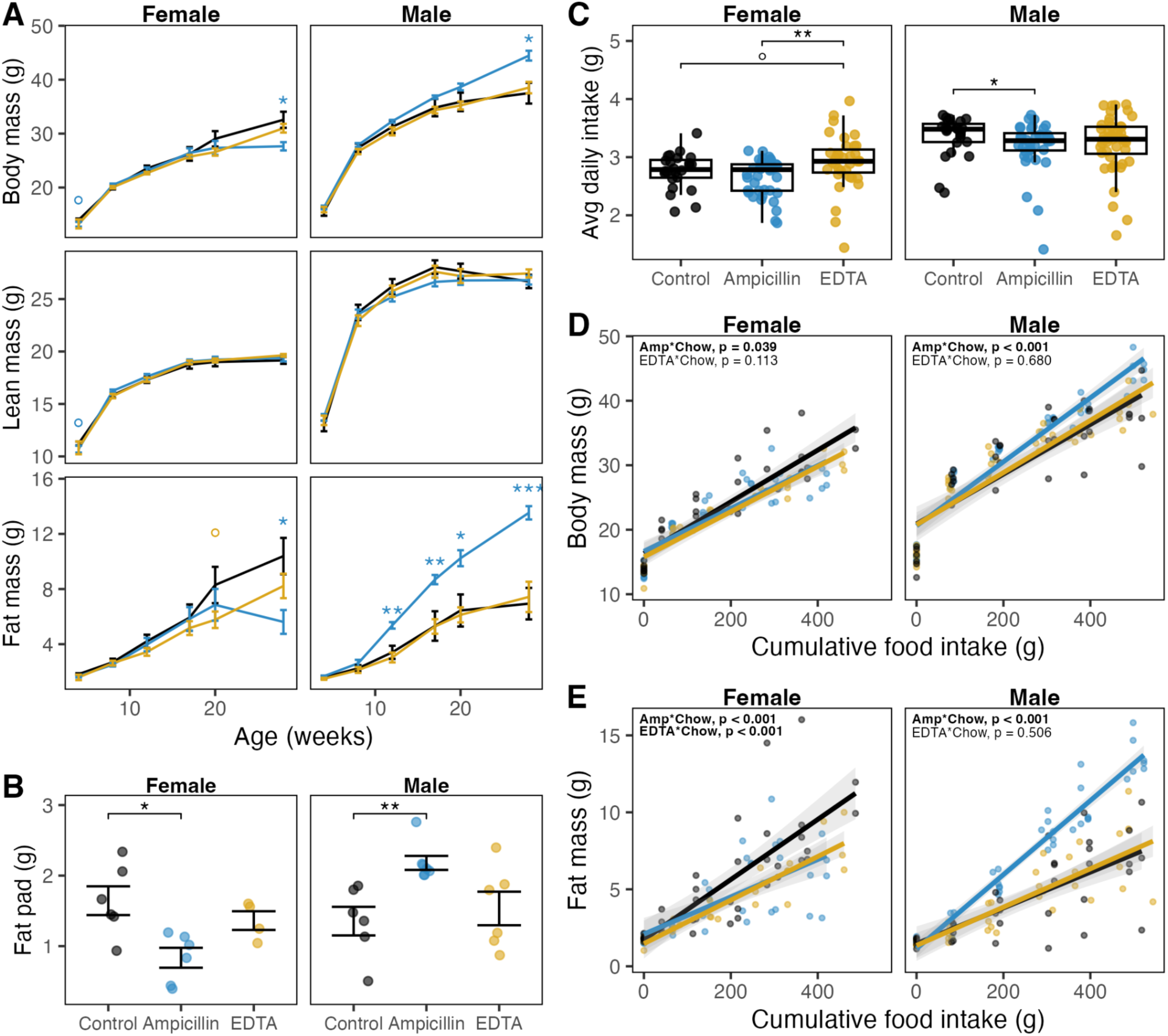
Growth, body composition, and food intake of mice in the long-term trial. (A) Body mass of mice from 4 through 28 weeks of age, with lean and fat body mass indexed by EchoMRI. (B) Mass of the gonadal white adipose tissue deposits at 28 weeks for females (parametrial fat pad) and males (epididymal fat pad). (C) Average daily food intake from weeks 4 through 28. Boxplots indicate median, first and third quartiles, with whiskers indicating 1.5 times interquartile range. (D-E) Total body mass (D) and body fat (E) as a function of cumulative food intake from 4 to 28 weeks of age, notated with results of linear mixed effects model ∼ Cumulative chow * Treatment with mouse ID as a random effect. Data are mean ± SE (A, B). All statistical annotation is treatment relative to the control group, unless otherwise indicated. Wilcoxon rank-sum test, ° = p<0.1, * = p<0.05, ** = p<0.01, *** = p<0.001.

Sex-specific effects of early-life antimicrobial treatment were even more pronounced after accounting for differences in food intake, as ampicillin-treated males exhibited lower food intake and EDTA-treated females exhibited higher food intake than sex-matched untreated controls (Figure 4C). To account for these differences in daily food intake, we used linear mixed effects models to evaluate how body mass and fat mass changed as a function of food intake. These models revealed that ampicillin-treated males gained markedly more weight and body fat per gram of food intake, and that EDTA– and ampicillin-treated females gained markedly less weight and particularly body fat per gram of food intake than sex-matched untreated controls (Figure 8D-E, p<0.05, LME).

Since differences in body mass and body composition among treatment groups could not be attributed to differences in food intake, we reasoned that they must result from either differential energy absorption from food or differential energy expenditure. To estimate unabsorbed energy, we first collected the feces produced by each mouse over 24 hours and measured fecal energetic density via bomb calorimetry. EDTA treatment led to higher fecal energy density in both female and male mice (Figure S7A). However, high variability in total 24-hour fecal production among EDTA-treated mice meant that the increased fecal energy density did not necessarily translate to higher total energy excretion (Figure S7B-C). Among ampicillin-treated mice, only males exhibited any differences in fecal energy excretion patterns, with higher fecal energy density but lower daily fecal production contributing to a non-significant net trend of modestly lower total energy excretion (p=0.268, Wilcoxon rank-sum test), about 14.4% lower than sex-matched untreated controls. Augmented energy harvest by ampicillin-treated males may have contributed in part to their increased weight and adiposity per gram of food intake.

To examine how the gut microbiota might independently contribute to altered host energy harvest, we used gas chromatography-mass spectrometry (GC-MS) to quantify cecal short-chain fatty acids (SCFAs), the major products of carbohydrate fermentation by the gut microbiota that serve as signaling molecules and metabolic fuel for diverse host tissues^21^ (Figure S8). For nearly all SCFA types, there was a consistent effect of sex detected across treatment groups in a 2-way ANOVA, with higher SCFA concentrations observed in females. This surprising result may indicate either that a greater fraction of dietary nutrients enters the cecum in females, thus indicating lower ileal digestibility, or that female and male gut microbiota have differential fermentation capacity. Including sex as a covariate in follow-up testing, ampicillin-treated mice had higher concentrations of cecal propionate (p=0.035, Tukey’s HSD), valerate (p=0.032), isovalerate (p=0.067), and isobutyrate (p=0.082) compared with untreated controls, while EDTA treatment was associated with a marginally significant reduction in acetate (p=0.053) and total SCFAs (p=0.068), with the latter result driven largely by acetate as the most abundant SCFA. Given phenotypic sex differences, we also included a sex by treatment interaction term in our tests of cecal SCFAs, which revealed borderline significant increases in total SCFAs (p=0.058), propionate (p=0.061), and butyrate (p=0.062) in ampicillin-treated females versus males.

Jointly, these data suggest that while differential dietary energy harvest likely contributed to the increased body mass and adiposity observed in ampicillin-treated males, it only partially explains the decreased body mass and adiposity observed in EDTA– and ampicillin-treated females. We therefore tested whether these phenotypes might additionally be driven by differences in energy expenditure.

We used indirect calorimetry to estimate the resting energy expenditure (REE) of treated and untreated mice. Consistent with their lean phenotypes, ampicillin-treated females had higher body mass-corrected REE than untreated controls, with EDTA-treated females intermediate between these groups (Figure S9B), but REE did not significantly vary by treatment for females on an absolute basis, when corrected by lean mass only, or after when corrected for both lean mass and fat mass independently using ANCOVA (Figure S9). Similarly, consistent with their increased stores of inexpensive body fat without reduction in expensive lean mass, ampicillin-treated males exhibited similar REE to controls and EDTA-treated males on an absolute basis or when correcting for lean mass alone (Figure S9). Interestingly, correcting for both lean mass and fat mass using ANCOVA suggested a substantial ∼19% reduction in the REE of ampicillin-treated males, although this analysis was underpowered and did not reach statistical significance (p=0.149). Regardless, it has been noted previously that even very small differences in energy expenditure – e.g., 3-5% differences that are hard to detect without sample sizes on the order of n=100 – can meaningfully contribute to differential weight gain and body composition^22^.

Given the potential for even slight differences in REE to contribute to body composition outcomes, we next considered the different tissues that may have underpinned a more or less energetically costly body, as most internal organs have higher mass-specific metabolic rates than do muscles at rest^23^. While EDTA-treated mice showed few significant differences from controls, ampicillin treatment led to striking sex-specific effects on organ size that were broadly consistent with observed treatment-induced trends in body composition. Ampicillin-treated females exhibited lower combined internal organ masses compared with controls (Figure S10I). While this result was driven primarily by their smaller livers (Figure S10C), ampicillin-treated females also displayed marked increases in brain size (Figure S10H), a tissue with a high mass-specific metabolic rate^24^. In contrast, ampicillin-treated males exhibited higher combined masses of metabolically expensive organs compared with controls (Figure S10) and differences in organ sizes that were generally in the opposite direction to those seen in ampicillin-treated females, including larger livers, longer small intestines, and a non-significant trend of longer large intestines. These increased structures for digestion may underpin the higher energy harvest observed for ampicillin-treated males.

Taken together, our analyses of energy harvest, energy expenditure, and tissue allocation suggest that ampicillin-induced increases in body fat in males may arise in part from increases in energy absorption from food (potentially promoted by larger livers and longer intestines) coupled with overall conservation of resting energy expenditure. By contrast, decreased adiposity in ampicillin-treated and EDTA-treated females may arise in part from energy expenditure driven by unexpected increases in brain size in ampicillin-treated females and modest reductions in cecal acetate and energy absorption from food by EDTA-treated females.

## DISCUSSION

We aimed to test whether consumption of dietary preservatives, both traditional and industrial, might perturb the gut microbiota—an important contributor to human energy budgets. We found that physiologically relevant concentrations of common dietary preservatives affected the gut microbiota *in vitro*, *ex vivo*, and in live mice. Each preservative left a unique signature on the gut microbiota that was distinct from that of antibiotics. We further showed that early-life treatment with low levels of EDTA reduces energy absorption from food and fat storage in females but not males. Importantly, we tested only low levels of these preservatives that are within diet-relevant ranges, underscoring the possibility that such consumption of dietary preservatives might have similar effects in humans.

Different preservatives left unique signatures on gut microbiota composition both *ex vivo* and *in vivo*. This is consistent with previous work that found many non-antibiotic drugs to have antimicrobial properties against gut bacteria^13^ and recent evidence that non-antibiotic drugs have mechanisms of action that are highly diverse and largely distinct from those of antibiotics^25^. Differing mechanisms of host absorption, metabolism, and excretion may also account for the varied effects of each preservative on the gut microbiome. Compounds may remain active once absorbed, but be excreted through a pathway (such as urine) that minimizes contact with the gut microbiota. Alternatively, a compound may remain unabsorbed and reach the distal gut but be inactivated or functionally altered by host or microbial metabolism, as is the case with bile acids and a number of drugs^14^. The effects of preservatives on whole community composition were generally stronger and broader *ex vivo* than *in vivo*, affecting taxonomic composition from phylum-level through ASV-level *ex vivo* but mainly affecting family-level through ASV-level composition within the mouse gut. These *ex vivo* versus *in vivo* differences are unsurprising as, in order to deliver biologically relevant doses of preservatives to mice (based on FDA acceptible daily intake limits), preservative concentrations in the mouse drinking water were only 10-20% of FDA-permitted maximum concentrations in food, meaning that administered compound concentrations were lower *in vivo* than in the *ex vivo* culture media.

We initially hypothesized that early-life treatment with preservatives would alter the gut microbiome and host metabolism in a manner similar to that of subtherapeutic antibiotics, ultimately inducing greater body fat in adults, especially males^26^. Our experiments with ampicillin confirmed these sex differences and offer novel insight into the energetic basis of early-life antibiotic-induced adult adiposity in males, in which higher dietary energy harvest (possibly due to longer small intestines) and marginally lower resting metabolic rate conspire to induce positive energy balance. In females, ampicillin did not promote but rather reduced adiposity. Exposure to EDTA also led to significant reductions in body fat in treated females compared with untreated female controls, an effect driven by lower dietary energy harvest. These antimicrobial-induced reductions in female energy status have not been identified previously, to our knowledge, potentially because leanness has been a lesser focus than obesity among most researchers rooted in industrialized contexts.

We were particularly intrigued by the impact of antimicrobial compounds and host sex on cecal SCFAs, as widespread evidence links SCFA production by gut microbes to regulation of energy metabolism and fat storage^1^. The generally higher levels of all SCFAs we observed in females compared to males have not been reported elsewhere, to our knowledge, but may contribute to the observed sex differences in body composition. As expected, our subtherapeutic dosing of ampicillin increased the abundance of some SCFAs, notably propionate. On the other hand, EDTA treatment reduced acetate and total SCFAs, suggesting that EDTA-exposed gut microbiomes may be less efficient at fermenting carbohydrates reaching the colon, thus contributing to higher caloric contents in feces.

Importantly, EDTA was capable of altering the gut microbiota in early life, a critical period in which gut microbiome disruption can affect lifelong metabolic programming^16^ and immune development, although the latter was not the focus of this study. Further studies are necessary to understand how EDTA and other preservatives may affect the gut microbiome and host metabolism in humans, but our findings that EDTA treatment can inhibit nutrient absorption and adiposity in female mice compared to untreated sex-matched controls raises concerns about the use of currently permissible levels of EDTA in foods available to young children and animals. EDTA has many uses beyond food preservation, with applications in medicine, as an undisclosed component of other food additives (e.g., some artificial sweeteners contain EDTA as a stabilizer), and even as a vehicle for iron supplements added to breakfast cereals or given to children to prevent iron deficiency^27,28^. Future work to elucidate how preservatives and other xenobiotic compounds interact with the gut microbiota, and potential downstream consequences for host physiology, will be critical in identifying the ecological levers at our disposal for modulating metabolic health via the gut microbiome.

## METHODS

### Compound concentration calculations

Based on prior literature showing antimicrobial effects *in vitro* against foodborne pathogens, we screened 9 common food preservation agents, 7 synthetic compounds [BHA (butylated hydroxyanisole), BHT (butylated hydroxytoluene), disodium EDTA (ethylenediaminetetraacetic acid), sodium benzoate, sodium nitrate, sodium sulfite, sulfur dioxide] and 2 more traditional compounds [acetic acid (vinegar) and sodium chloride (table salt)] (Table S1). We selected these compounds due to their widespread use and because they represent main classes of preservatives in general use today.

For *in vitro* and *ex vivo* experiments, we tested the growth effects of each compound at concentrations of 20-2000 µg/ml, a range that includes the FDA maximum concentration for all regulated compounds. For *in vivo* mouse experiments, drinking water doses of each compound were calculated as follows: acceptable daily intake (ADI) as stated by the FDA was scaled allometrically to mice by a factor of 12.3, based on the relative body-mass-to-body-surface-area ratios of humans (37 kg/m^2^) and mice (3 kg/m^2^)^29^. Finally, we assumed an average daily water intake of 0.15 ml/g body mass for mice^15^, which was ultimately consistent with measured water intake (Figure S3), resulting in drinking water concentrations of 10% v/v (acetic acid), 41 mg/L (BHA), 206 mg/L (EDTA), and 57 mg/L (sodium sulfite). Ampicillin was administered at a concentration of 6.7 mg/L, a value modeled on previous studies^15^.

Commercial preparations of acetic acid (A1009), BHA (BH104), BHT (B1095), EDTA (E1001), sodium benzoate (S1146), sodium nitrate (SO183), and sodium sulfite (S1113) were obtained from Spectrum Chemicals (New Brunswick, NJ); sulfur dioxide (sc-215934) from Santa Cruz Biotechnology (Santa Cruz, CA); ampicillin trihydrate (J66514) from Thermo Fisher (Waltham, MA); and sodium chloride (BDH9286) from VWR International (Radnor, PA).

### Growth of bacterial strains and whole communities

All *in vitro* and *ex vivo* culturing was performed under anaerobic conditions using Brain Heart Infusion (BHI) broth (BD 214010) supplemented with yeast extract (5 g/L, VWR J850) and resazurin sodium salt (0.1 mg/L, Sigma Aldrich R7017) that was autoclaved before the final addition of L-cysteine hydrochloride (0.5 g/L, Sigma Aldrich C7477). Media and all consumables were allowed to reduce in an anaerobic chamber for >12 hours before use. All tests were performed in optically clear, flat-bottomed 96-well plates filled with 190 µl broth pre-mixed with the appropriate concentration of each compound with 10 µl inoculum. For tests of individual strains, the inoculum consisted of *Bacteroides ovatus* (ATCC 8483), *Clostridium symbiosum* (ATCC 14940), *Escherichia coli* (ATCC 47076), or *Eggerthella lenta* (ATCC 25559) grown to mid-logarithmic growth phase and normalized to an optical density at 600 nm (OD_600_) of 0.1. For whole fecal communities, fecal samples were collected fresh from adult C57BL6/J mice and moved to an anaerobic chamber within 10 minutes of collection. Samples were then diluted 1:30 in pre-reduced PBS, vortexed for 10 minutes to homogenize, and the resulting cell suspension was used as the inoculum. Plates were incubated at 37°C and all combinations of inoculum/compound/concentration were performed in triplicate.

We validated OD_600_ readings of cell density with counts of colony forming units (CFUs) in for a random selection samples, cultured on agar plates (Figure S2), made from the supplemented BHI media described above plus 15g/L agar (BD 214010). Cultures were diluted 10-fold 8 times to generate dilutions from 1:10 through 1:10^8^. We plated 4 5µl replicates of each dilution and incubated these in an anaerobic chamber for 24-48 hours, until colonies were visible (example image: Figure S2D). CFU counts of cell density were generally well correlated with OD_600_ readings.

### Animal husbandry

For the 7-day trial, 5 groups of 6 male, 7-week-old C57BL/6J cage-mate mice were purchased from Jackson Laboratory. Mice were transferred to individual housing shortly after arrival and were maintained in separate cages for the duration of the experiment. To prevent baseline variation in the gut microbiome among source cages from biasing results, we randomly assigned one mouse from each cage-mate group to one of 6 treatment groups: water (negative control), ampicillin (positive control), acetic acid, BHA, EDTA, or sodium sulfite. After the 7^th^ day of treatment, mice were sacrificed by CO2 inhalation and samples of gut effluent collected.

For the long-term, developmental experiment, 9 timed-pregnant mice at 11 days gestation were delivered from Jackson Laboratory, with n=3 assigned to each treatment group (control, EDTA, ampicillin). At 13.5 days gestation, mice in the EDTA and ampicillin groups were given treated drinking water. Drinking water was refreshed twice weekly and treatment continued for the duration of the experiment, until offspring were 28 weeks old. Offspring were weaned at 3 weeks of age, and housed together by litter and sex, with ≤5 mice per cage. Starting at 4 weeks of age and every 4 weeks thereafter, we assessed mouse body composition via EchoMRI and collected fecal samples for gut microbial profiling. At 28 weeks, mice were sacrificed, and samples of gut effluent and other tissues were collected for downstream analysis.

Mice in both experiments were fed irradiated PicoLab Mouse Diet 20 5058 provided *ad libitum*. All mouse experiments were performed in the specific pathogen-free Harvard University Biological Research Infrastructure facility under a protocol approved by the Harvard University Institutional Animal Care and Use Committee (Protocol #17-06-306).

### Gut microbial profiling via 16S rDNA sequencing

We assessed gut microbial community composition of bacterial cultures, feces and gut effluent via 16S rDNA sequencing. For the long-term treatment experiment where mice were housed in groups, one mouse from each cage was randomly selected for gut microbial profiling, as it is generally inappropriate to treat co-housed animals as independent biological replicates for the purposes of gut microbiome profiling due to coprophagy and other sources of extensive horizontal transmission. We isolated DNA using the Qiagen Powersoil DNA Isolation Kit following manufacturer’s instructions. Next, we performed PCR amplification of the 16S rRNA gene using custom-barcoded 515F and 806R primers targeting the V4 region of the gene. We performed PCR on each sample in triplicate with sample-specific negative controls with the following protocol: 95°C for 3 min; 35 cycles of 94°C for 45 s, 50°C for 30 s, and 72°C for 90 s; and 10-minute final extension at 72°C. We then cleaned amplicons with AmpureXP beads (Agencourt) and quantified samples with Quant-iT Picogreen dsDNA Assay Kit (Invitrogen) prior to pooling samples evenly by DNA content. The resulting 16S rDNA libraries underwent 1×150 bp sequencing across 3 lanes of an Illumina HiSeq, with one lane dedicated to each of the *in vitro* plus *ex vivo* samples, 7-day mouse study samples, and long-term mouse study samples.

Sequences were processed in QIIME2^30^, first by de-noising with Dada2 and truncating at 149 bp to ensure maximum sequence quality, resulting in read depths of 126,218 ± 25,902 (*in vitro* and *ex vivo* study), 172,173 ± 37,136 (7-day study), and 68,271 ± 10,532 (long-term study). Taxonomy was assigned using the GreenGenes classifier^31^ and a rooted tree for all amplicon sequence variants (ASVs) was generated. The taxonomy, phylogeny, and ASV feature table were then imported into R (version 4.3.2) using qiime2R (version 0.99.6). Pre-processing was conducted using phyloseq (version 1.46.0^32^). First, each sample was pruned of very low abundance ASVs, defined as ≤3 reads per study pool. Next, reads were subsampled evenly at 50,000 reads/samples for the *in vitro* plus *ex vivo* and 7-day *in vivo* studies, and at 40,000 reads/sample for the long-term study, which excluded 2 samples with <40,000 reads, resulting in n=124, n=327, and n=182 samples in each study, respectively. Further processing of 16S rDNA sequences was then performed in R using phyloseq for calculating distance matrices and ordinations; vegan (version 2.6-4) for PERMANOVA tests; and MaAsLin2 (version 1.16.0) for identifying differentially abundant taxa using general linear models.

### Quantitative PCR (qPCR) of 16S rRNA gene

We performed qPCR on the V4 region of the 16S gene (515F and 806R, non-barcoded primers) in triplicate, using a standard curve on each plate based on genomic DNA isolated from a pure culture of *Escherichia coli*. We used the following recipe for each PCR reaction: 12.5 μl SYBR Green qPCR mix, 2.25 μl of each non-barcoded primer (515F and 806R), 6 μl nuclease-free H2O, and 2 μl template DNA, for a total volume of 25 μl per well. We ran the qPCR reaction in a BioRad CFX 96-well Real-Time PCR thermocycler with the following protocol: initial denature at 94°C for 15 min; 40 cycles of 95°C for 15 s, 50°C for 40 s, and 72°C for 30 s. To calculate 16S rRNA gene abundance, we first multiplied DNA concentrations of each sample as measured via qPCR then divided by the mass of the original fecal sample and multiplied by 2.03 x 10^5^—an estimate of genome-equivalents per ng DNA based on a mean gut microbial community genome size of 4.50 Mbp^11^.

### Resting energy expenditure

We used an open-flow indirect calorimetry system for measurement of resting energy expenditure, using Classic Line instrumentation manufactured by Sable Systems International (Las Vegas, NV), as described previously^33^.

Mice were fasted for 4 hours before being placed in respirometer chambers and given 1 hour to acclimate to the chambers before measurement began. Measurements spanned 1 hour, with continuous activity measurement via force plates. Gas flow into the oxygen analyzer cycled between one of 2 mouse cages every 7 minutes, with a 7-minute baseline between each cycle. Raw data was processed in ExpeData3, where flow rates were corrected for standard temperature and pressure (STP) and O_2_ readings were corrected by spanning dry baseline air to 20.95% O_2_. Oxygen consumption (VO_2_) (ml/min) was then calculated as:

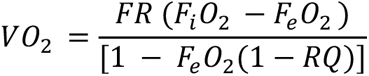

Where *FR* is the dry flow rate, *F_i_0_2_* is the % oxygen in incurrent (baseline) air, *F_e_0_2_* is the % oxygen in excurrent (post-chamber) air, and RQ is the respiratory quotient, here assumed to be 0.8^34^. Oxygen consumption was converted to energy expenditure using the oxyjoule equivalent of 20.13 J/ml O_2_^34^, then further converted into kcal/day by a factor of 4184 J/kcal.

To find resting energy expenditure (REE) from the continuous measurement of energy expenditure (EE), we first averaged EE over 20 second increments, then found the 3 minimum EE values ≥1 minute apart with confirmed minimal activity, then averaged these 3 values as the estimated REE. Since researchers have variously championed the biological relevance of uncorrected and corrected REE values^22,35^, we have elected to report absolute REE as well as REE corrected for body mass, lean mass, or both via ANCOVA.

### Estimation of fecal energy excretion

In the long-term *in vivo* study, all mice were housed individually in fresh cages for 24 hours after which the bedding was collected and sifted for feces. Collected feces were desiccated by freeze drying to calculate the total dry-weight fecal production per day. The entire sample of collected feces was then was then combusted in a bomb calorimeter (Parr Instrument Co., 6050 Calorimeter).

### Statistical analysis

For data not obtained via sequencing, we performed all statistical analysis within the R Studio platform (version 023.09.1+494) and the tidyverse packages (version 2.0.0). For single-variate, non-normally distributed comparisons between treatment groups, we performed Kruskal-Wallis tests, followed by pairwise Wilcoxon rank-sum tests with the control as a reference group. For normally distributed data, we performed ANOVAs followed by Tukey’s HSD. For longitudinal data where individual mice were sampled multiple times, we used linear mixed effects models to control for the random effects of each mouse and avoid autocorrelation, with models run using the nlme package (version 3.1-164). Multivariate analyses of microbiome composition were performed on sample distance matrices using the PERMANOVA test in the vegan package.

## DATA AVAILABILITY

Sequencing data have been deposited to the NCBI Sequence Read Archive under submission number SUB14437337.

## AUTHOR CONTRIBUTIONS

The project was conceived and designed by LDS and RNC. Culturing experiments were performed by LDS and CRAB. Animal work was performed by LDS and KSC. Samples were processed for sequencing by LDS and EMV. All analysis and other sample processing was performed by LDS. The project was supervised by RNC. The manuscript was drafted by LDS and RNC and revised with feedback from all authors.

## ACKNOWLEDGMENTS

We thank Martin Blaser and members of the Carmody lab for helpful discussions and feedback on early drafts of this manuscript. This study was supported by a National Science Foundation Graduate Research Fellowship (to LDS) and awards from The William F. Milton Fund (to RNC), Harvard Dean’s Competitive Fund for Promising Scholarship (to RNC), and National Science Foundation (BCS-2142073 to LDS and RNC).

## SUPPLEMENTARY MATERIALS

**Figure S1.**
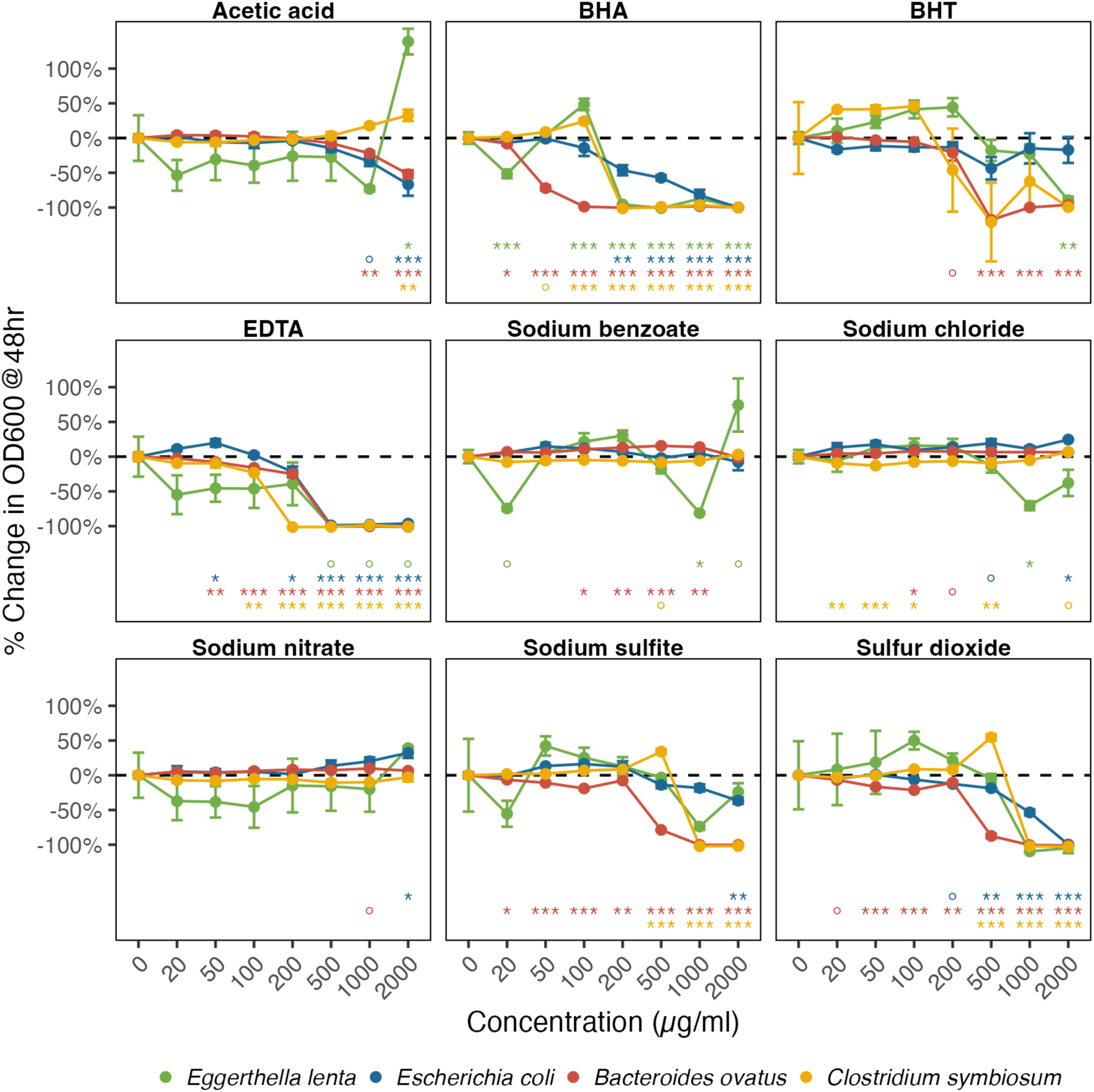
Impact of preservatives on growth of individual gut isolates *in vitro*. Cell density (OD_600_) for each preservative and concentration is shown as a percent of no-compound growth controls after 48 hours growth. Data represent mean ± SE of 3 technical replicates. Statistical differences for each strain at each concentration versus no-compound controls (0 µg/ml) using Tukey’s HSD test annotated as: ° = p<0.1, * = p<0.05, ** = p<0.01, *** = p<0.001.

**Figure S2.**
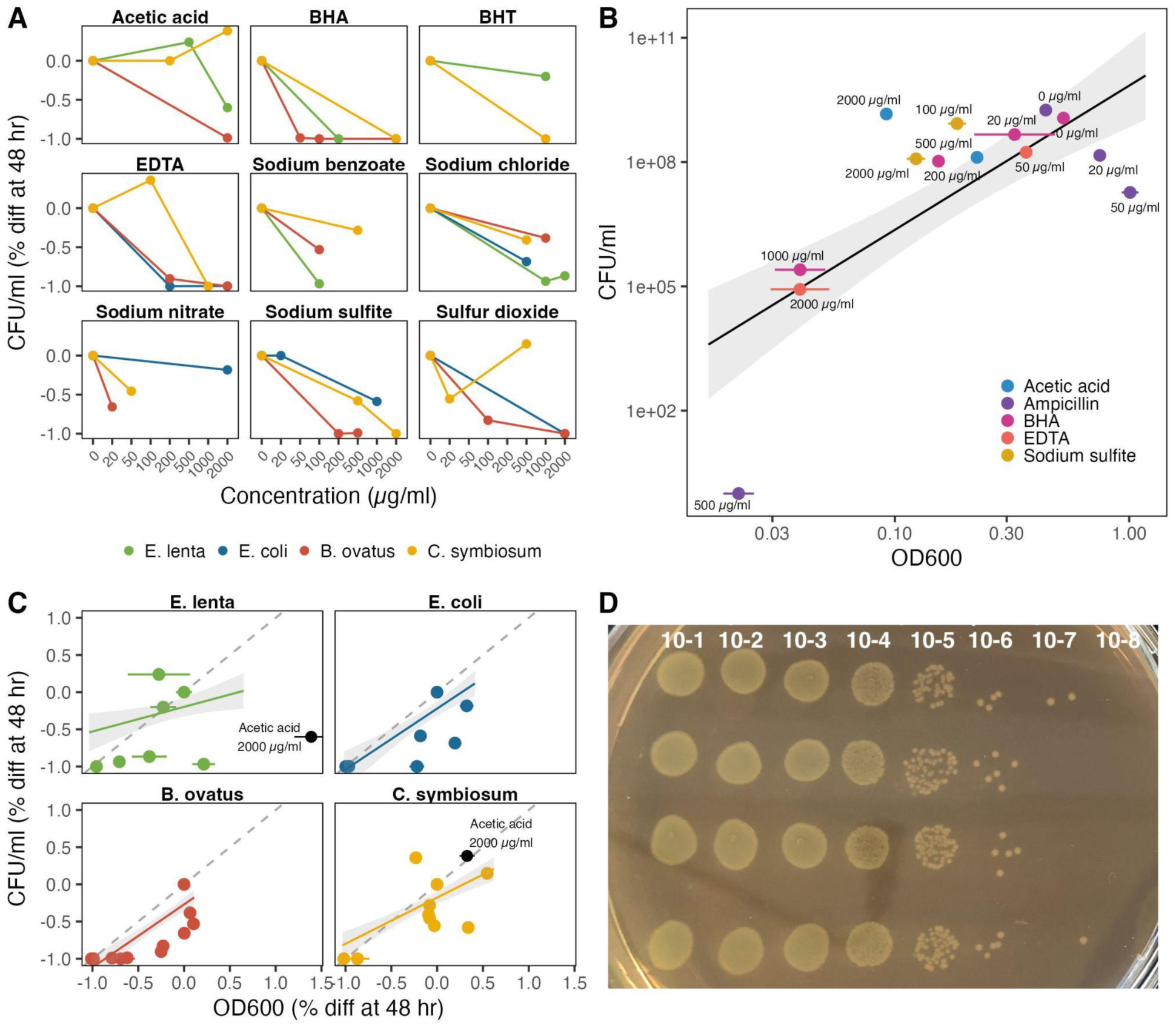
Validation of bacterial cell density measurements. Counts of colony forming units (CFUs) from a subset of *in vitro* cultures of both individual strains expressed as the percent difference from controls at endpoint (A, C) and whole fecal communities at endpoint (B, all expressed relative to optical density readings). Data in B and C are mean ± SE, with panel B plotted on a log-log scale to highlight CFU counts on the lower end of the spectrum. (D) Image of a representative test plate, with 4 replicates of each 5 µl drop at each dilution (1:10^1^ through 1:10^8^).

**Figure S3.**
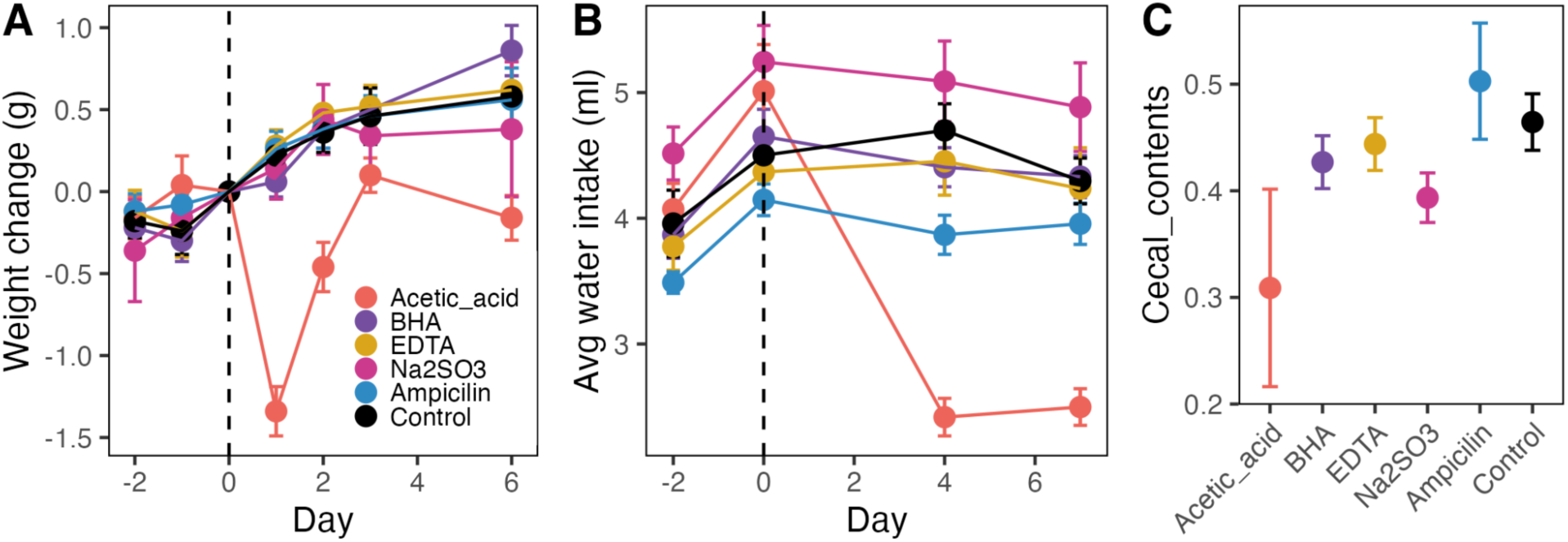
Physical measurements in 7-day *in vivo* trial. (A) Change in mouse body mass from Day –2 (Baseline) through Day 6 of treatment. (B) Water intake during baseline and treatment. Values for each day represent the change in mass of cage water bottles divided by the days since the last measurement. (B) Mass of cecal contents at Day 7. Data are mean ± SE.

**Figure S4.**
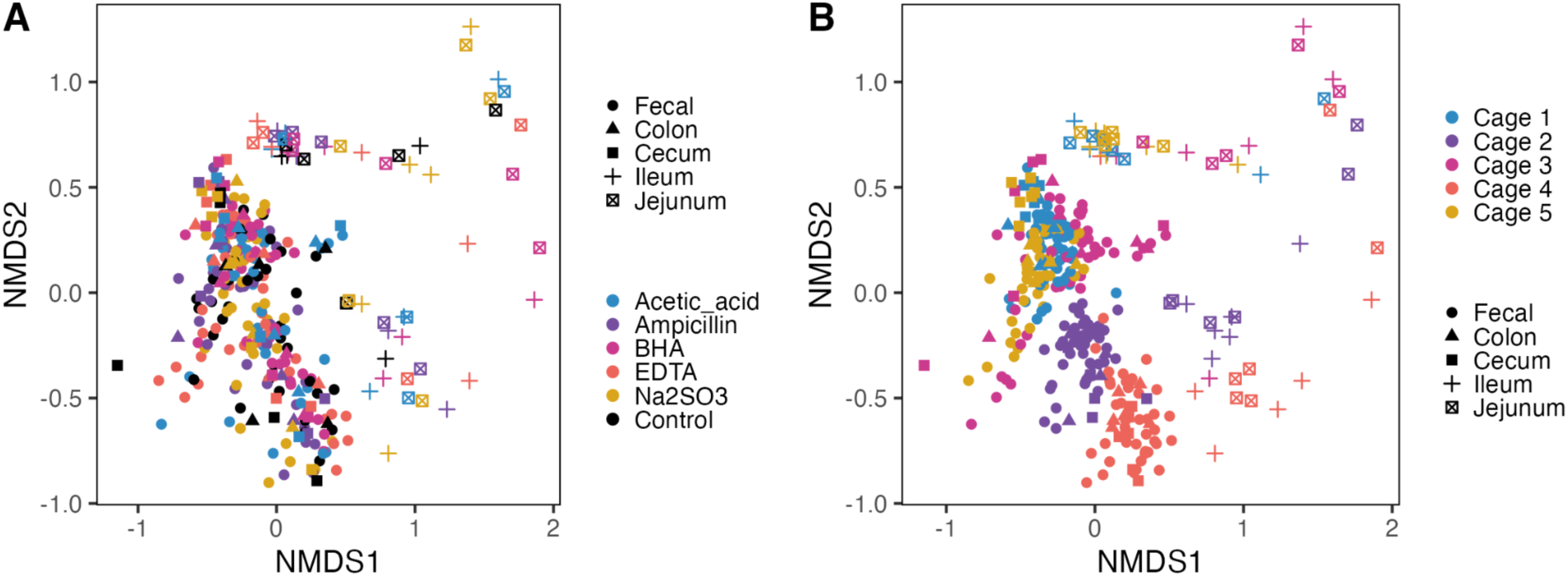
Gut microbial community composition in 7-day *in vivo* trial. Non-dimensional linear scaling (NMDS) ordination plot of Bray-Curtis distances between gut microbiome samples collected along the gastrointestinal tract. (A) Samples colored by treatment group. (B) Samples colored by pre-baseline cage groupings to emphasize cage effect.

**Figure S5.**
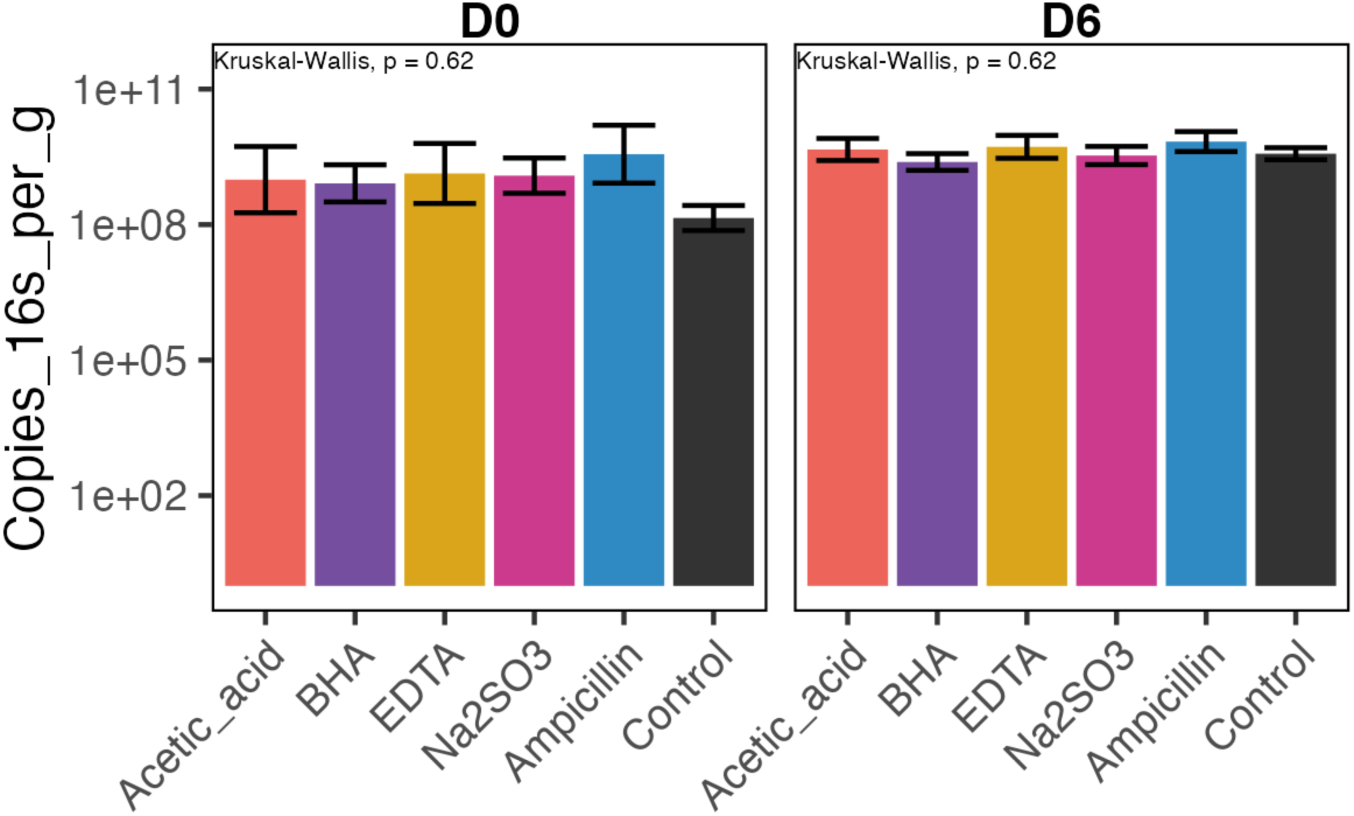
Absolute fecal bacterial abundance at Day 0 and Day 6, measured by quantitative PCR of the 16S gene. Data are mean ± SE.

**Figure S6.**
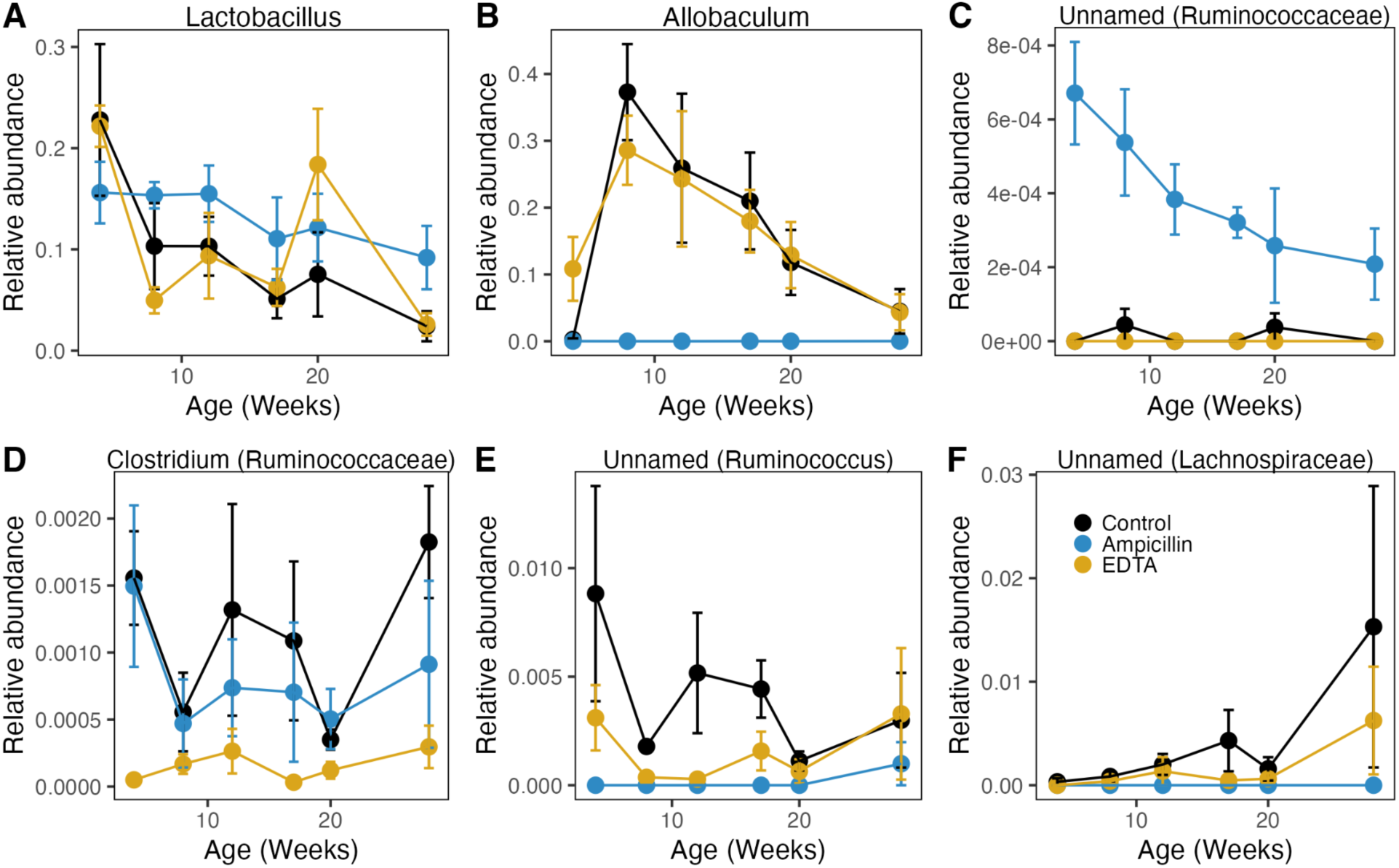
Relative abundance of select genera and ASVs in the long-term trial. Notable genera previously found to differ with early-life antibiotic treatment^16^ (A, B) and genera and unnamed ASVs identified by MaaAslin2 as significantly different between at least one treatment group and controls (C-F). Across these taxa, we observed significant differential abundance by ampicillin treatment (B, C, E, F), by EDTA treatment (D-F), and by age (C, F). ASVs are presented with the lowest known taxonomic identification. Data are mean ± SE and colored by treatment group (n = 4-7).

**Figure S7.**
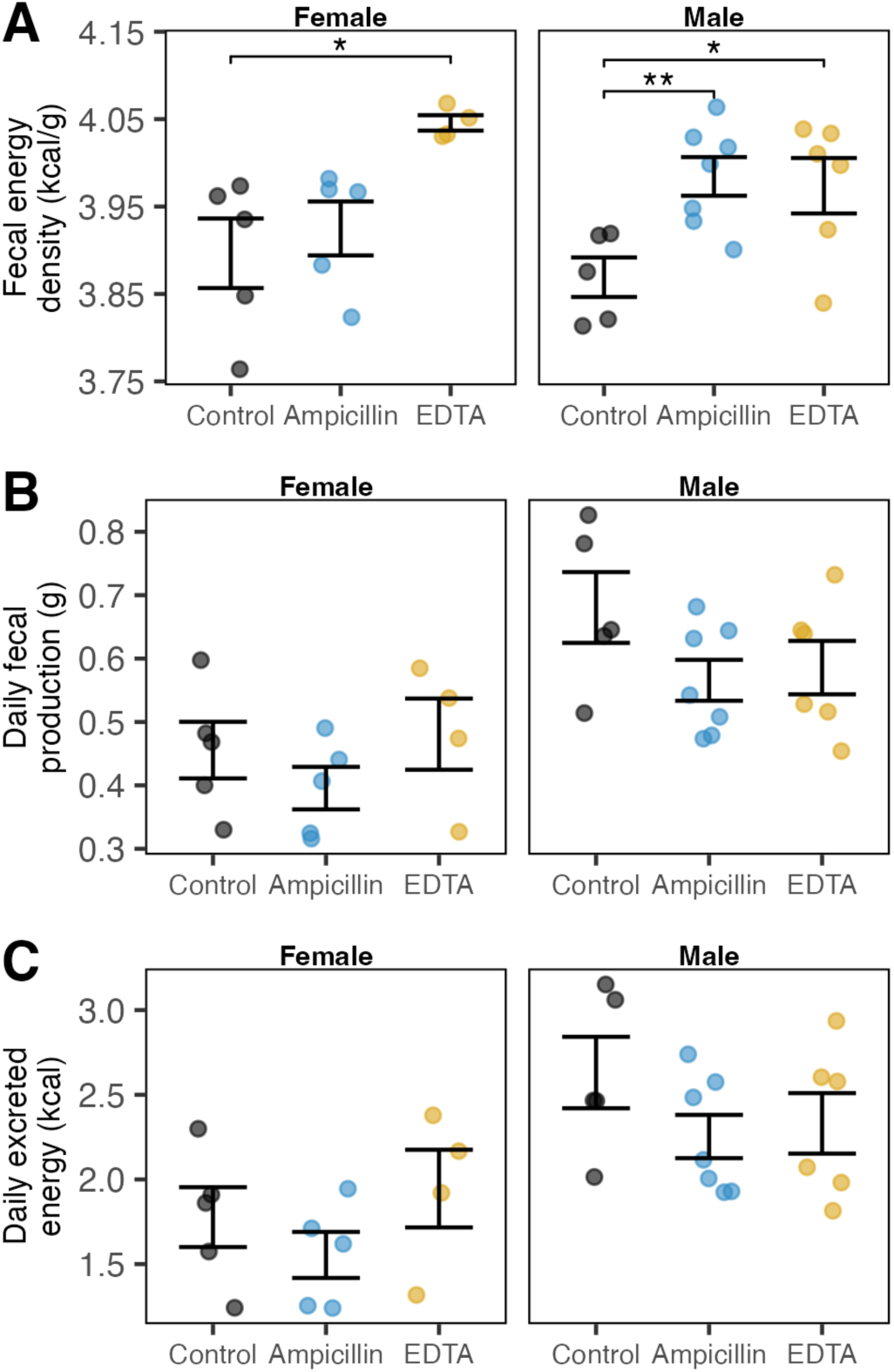
Fecal energy excretion. (A) Caloric density of fecal dry matter quantified via bomb calorimetry. (B) Total fecal production over 24 hours. (C) Total excreted energy over 24 hours, calculated as the product of fecal energy density and fecal production over the same time period. Material was collected from mice at 28 weeks of age. Error bars are mean ± SE. Wilcoxon rank-sum test, ° = p<0.1, * = p<0.05, ** = p<0.01, *** = p<0.001.

**Figure S8.**
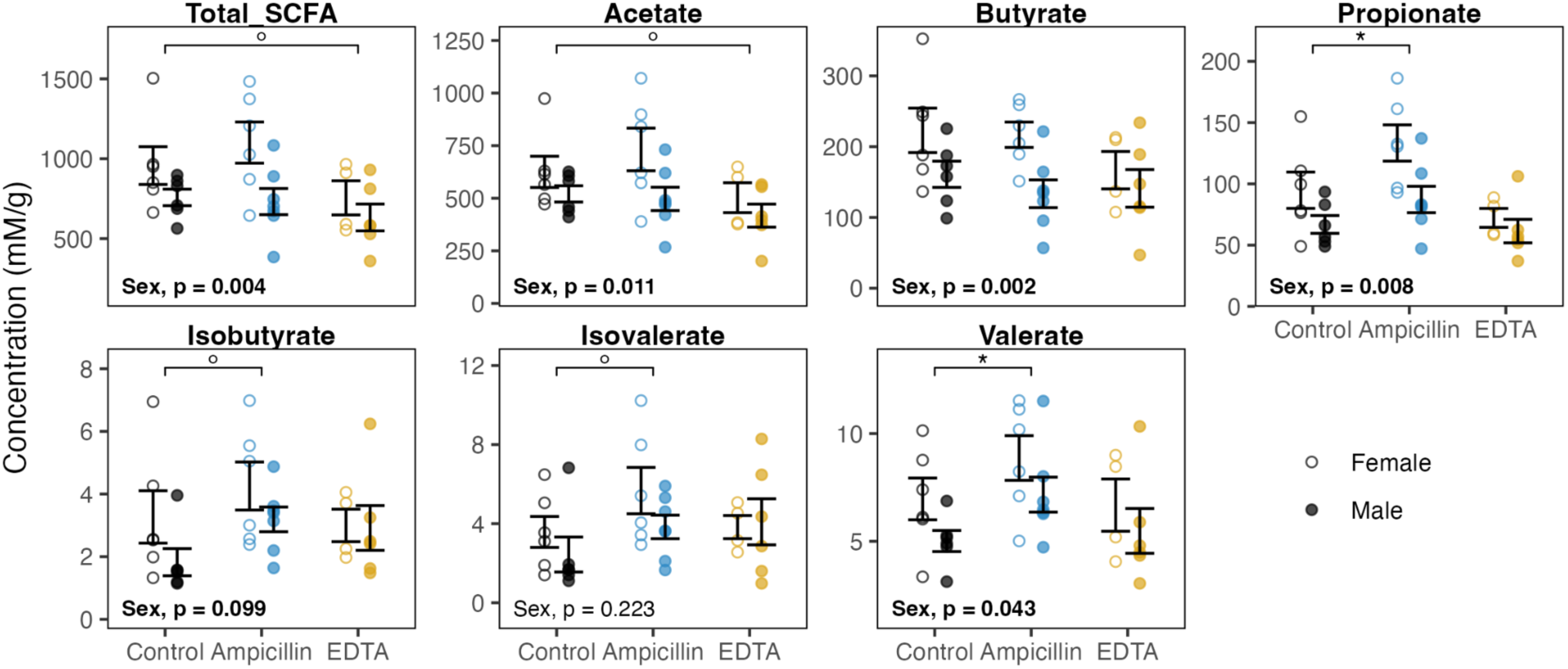
Cecal short-chain fatty acid contents. Concentrations measured by GC-MS and displayed as mM per g dry matter. Total SCFA is the sum of all 6 SCFAs shown individually. Data are mean ± SE. 2-way ANOVA on SCFA Concentration ∼ Sex * Treatment, followed by Tukey’s HSD, ° = p<0.1, * = p<0.05, ** = p<0.01, *** = p<0.001. Interaction effects: Total SCFAs (p=0.058), propionate (p=0.061), and butyrate (p=0.062) for ampicillin-treated females versus ampicillin-treated males.

**Figure S9.**
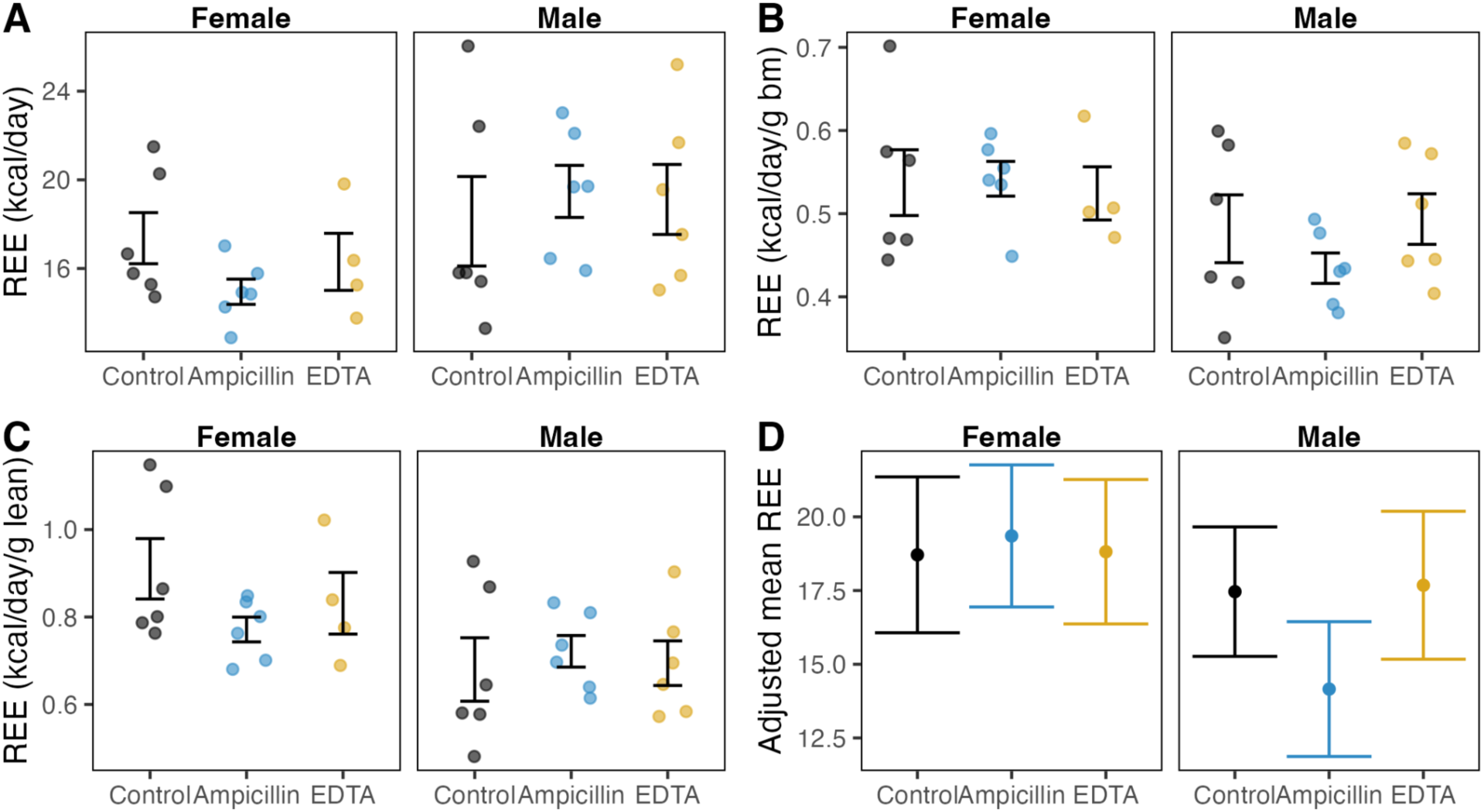
Indirect calorimetry. (A) Estimated resting energy expenditure (REE) of fasted 28-week-old mice, expressed as kcal per day. (B) REE expressed per gram of mouse body mass. (C) REE expressed per gram of lean body mass. Data in A-C are mean ± SE. No significant differences from control were detected for any group (Mann-Whitney U test, p>0.05). (D) REE group means adjusted using ANCOVA with mouse lean mass and fat mass as covariates. Data in D are adjusted means ± SE.

**Figure S10.**
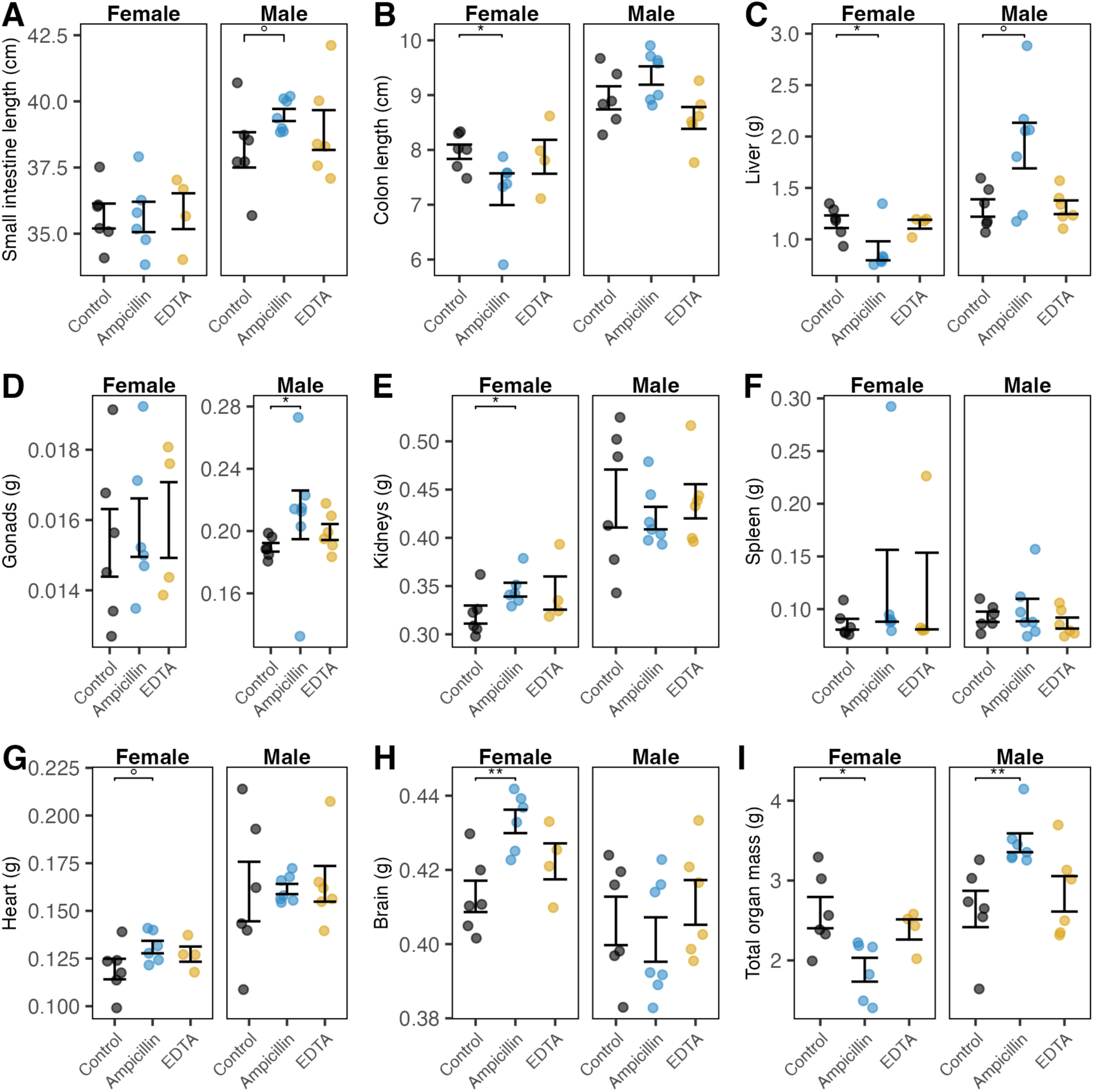
Organ sizes of 28-week-old mice. Organ sizes given as length (A, B) or mass (C-I). For paired organs (gonads and kidneys), value represents the sum of the left and right sides. Total organ mass (I) is the sum of all organ mass measurements (C-H). Data are mean ± SE. Wilcoxon rank-sum test, ° = p<0.1, * = p<0.05, ** = p<0.01, *** = p<0.001.

**Table S1:** Background information on preservatives tested in initial *in vitro* screen.

| Compound | Functional class | Food use | Acceptable daily intake† (mg/kg/day) | FDA concentration limits (µg/ml) |
| --- | --- | --- | --- | --- |
| Acetic acid | Antimicrobial, flavoring agent, acidity regulator | Vinegar (natural), cheese, dressings, condiments | NA | NA |
| BHA (butylated hydroxyanisole) | Antimicrobial | Cereals, chips, baked goods, condiments, shortening, fats/oils | 0.5<sup>36</sup> | 200 |
| BHT (butylated hydroxytoluene) | Antioxidant, antimicrobial | Same as BHA | 0.3<sup>37</sup> | 200 |
| Disodium EDTA (ethylenediamine-tetraacetic acid) | Antioxidant, antimicrobial<br>Antioxidant, color retention agent, preservative, sequestrant, stabilizer | Canned vegetables, potatoes, beans, sauces, dressings, sweeteners, multivitamins | 2.5<sup>38</sup> | 1000 |
| Sodium benzoate | Antimicrobial | Carbonated beverages, syrup, margarine, dressings;<br>Natural in cranberries, plums, cinnamon, cloves | 5<sup>39</sup> | 1000 |
| Sodium chloride | Flavoring agent | All food types, household use | NA | NA |
| Sodium nitrate | Antimicrobial | Processed meats | 3.7<sup>40</sup> | 500 |
| Sodium sulfite | Antimicrobial | Wine, cider, beer, cereal/potato-based snacks | 0.7<sup>41</sup> | 500<sup>§</sup> |
| Sulfur dioxide | Antimicrobial | Dried fruit, wine, cider, beer | 0.7<sup>41</sup> | 500 |

† Acceptable daily limit (ADI) is based on JECFA/WHO recommendations and represents the upper limit of the compound that is safe for ingestion in units of mg compound per kg body weight per day. Estimated dietary concentration (EDC) was based on FDA limits for compound concentrations in foods, when available. For acetic acid for which there is no usage limit, the upper dietary concentration was estimated based on dietary intake estimates.
§ While the FDA does not specify a limit of concentrations in food from our source, sodium sulfate is evaluated together with sulfur dioxide and the same limits are proposed by other regulatory bodies (namely FAO/WHO). Therefore we apply the permissible concentration of sulfur dioxide (0.05%) to sodium sulfite here.

**Tables S2 and S3:**

Please see appended Excel file.

## REFERENCES

1. Carmody, R. N. & Bisanz, J. E. Roles of the gut microbiome in weight management. Nat Rev Microbiol 21, 535–550 (2023).

2. Cani, P. D. et al. Microbial regulation of organismal energy homeostasis. Nat Metab 1, 34–46 (2019).

3. Chung, H. et al. Gut immune maturation depends on colonization with a host-specific microbiota. Cell 149, 1578–1593 (2012).

4. Osadchiy, V., Martin, C. R. & Mayer, E. A. The Gut-Brain Axis and the Microbiome: Mechanisms and Clinical Implications. Clin Gastroenterol Hepatol 17, 322–332 (2019).

5. Lynch, S. V. & Pedersen, O. The Human Intestinal Microbiome in Health and Disease. N Engl J Med 375, 2369–2379 (2016).

6. Turnbaugh, P. J., Bäckhed, F., Fulton, L. & Gordon, J. I. Diet-induced obesity is linked to marked but reversible alterations in the mouse distal gut microbiome. Cell Host Microbe 3, 213– 223 (2008).

7. David, L. A. et al. Diet rapidly and reproducibly alters the human gut microbiome. Nature 505, 559–563 (2014).

8. Carmody, R. N. et al. Cooking shapes the structure and function of the gut microbiome. Nat Microbiol 4, 2052–2063 (2019).

9. Caffrey, E. B., Sonnenburg, J. L. & Devkota, S. Our extended microbiome: The human-relevant metabolites and biology of fermented foods. Cell Metab 36, 684–701 (2024).

10. Suez, J. et al. Artificial sweeteners induce glucose intolerance by altering the gut microbiota. Nature 514, 181–186 (2014).

11. Roopchand, D. E. et al. Dietary Polyphenols Promote Growth of the Gut Bacterium Akkermansia muciniphila and Attenuate High-Fat Diet-Induced Metabolic Syndrome. Diabetes 64, 2847–2858 (2015).

12. Chassaing, B. et al. Dietary emulsifiers impact the mouse gut microbiota promoting colitis and metabolic syndrome. Nature 519, 92–96 (2015).

13. Maier, L. et al. Extensive impact of non-antibiotic drugs on human gut bacteria. Nature 555, 623–628 (2018).

14. Spanogiannopoulos, P., Bess, E. N., Carmody, R. N. & Turnbaugh, P. J. The microbial pharmacists within us: a metagenomic view of xenobiotic metabolism. Nat Rev Microbiol 14, 273–287 (2016).

15. Cho, I. et al. Antibiotics in early life alter the murine colonic microbiome and adiposity. Nature 488, 621–626 (2012).

16. Cox, L. M. et al. Altering the intestinal microbiota during a critical developmental window has lasting metabolic consequences. Cell 158, 705–721 (2014).

17. Chen, R.-A., et al. Dietary Exposure to Antibiotic Residues Facilitates Metabolic Disorder by Altering the Gut Microbiota and Bile Acid Composition. mSystems 7, e0017222 (2022).

18. Mallick, H. et al. Multivariable association discovery in population-scale meta-omics studies. PLoS Comput Biol 17, e1009442 (2021).

19. Takeuchi, T. et al. Gut microbial carbohydrate metabolism contributes to insulin resistance. Nature 621, 389–395 (2023).

20. Kimura, I. et al. Maternal gut microbiota in pregnancy influences offspring metabolic phenotype in mice. Science 367, eaaw8429 (2020).

21. den Besten, G., et al. The role of short-chain fatty acids in the interplay between diet, gut microbiota, and host energy metabolism. J Lipid Res 54, 2325–2340 (2013).

22. Speakman, J. R. Measuring energy metabolism in the mouse – theoretical, practical, and analytical considerations. Front Physiol 4, 34 (2013).

23. Wang, Z. et al. Specific metabolic rates of major organs and tissues across adulthood: evaluation by mechanistic model of resting energy expenditure. Am J Clin Nutr 92, 1369–1377 (2010).

24. Wheeler, P. E. An investigation of some aspects of the transition from ectothermic to endothermic metabolism in vertebrates. (Durham University, 1984).

25. Noto Guillen, M., Li, C., Rosener, B. & Mitchell, A. Antibacterial activity of nonantibiotics is orthogonal to standard antibiotics. Science 384, 93–100 (2024).

26. Cox, L. M. & Blaser, M. J. Antibiotics in early life and obesity. Nat Rev Endocrinol 11, 182–190 (2015).

27. Wreesmann, C. T. J. Reasons for raising the maximum acceptable daily intake of EDTA and the benefits for iron fortification of foods for children 6-24 months of age. Matern Child Nutr 10, 481– 495 (2014).

28. EFSA Panel on Food Additives and Nutrient Sources added to Food (ANS). Scientific Opinion on the use of ferric sodium EDTA as a source of iron added for nutritional purposes to foods for the general population (including food supplements) and to foods for particular nutritional uses. EFSA J. 8, 1414 (2010).

29. Reagan-Shaw, S., Nihal, M. & Ahmad, N. Dose translation from animal to human studies revisited. FASEB J. 22, 659–661 (2008).

30. Bolyen, E. et al. Reproducible, interactive, scalable and extensible microbiome data science using QIIME 2. Nat Biotechnol 37, 852–857 (2019).

31. DeSantis, T. Z. et al. Greengenes, a chimera-checked 16S rRNA gene database and workbench compatible with ARB. Appl Environ Microbiol 72, 5069–5072 (2006).

32. McMurdie, P. J. & Holmes, S. phyloseq: an R package for reproducible interactive analysis and graphics of microbiome census data. PLoS One 8, e61217 (2013).

33. Carmody, R. N. Energetic Consequences of Thermal and Non-Thermal Food Processing. (Harvard University, 2013).

34. Lighton, J. R. B. Measuring Metabolic Rates: A Manual for Scientists. (Oxford University Press, 2018).

35. Tschöp, M. H. et al. A guide to analysis of mouse energy metabolism. Nat Methods 9, 57–63 (2011).

36. Joint FAO/WHO Expert Committee on Food Additives. Evaluation of Certain Food Additives and Contaminants: Thirty-Third Report of the Joint FAO/WHO Expert Committee on Food Additives. (1989).

37. Joint FAO/WHO Expert Committee on Food Additives. Evaluation of Certain Food Additives and Contaminants: Fourty-Fourth Report of the Joint FAO/WHO Expert Committee on Food Additives. (1995).

38. Joint FAO/WHO Expert Committee on Food Additives. Toxicological Evaluation of Certain Food Additives with a Review of General Principles and Specifications: Seventeenth Report of the Joint FAO/WHO Expert Committee on Food Additives. (1973).

39. Joint FAO/WHO Expert Committee on Food Additives. Evaluation of Certain Food Additives and Contaminants. Thirty-Seventh Report of the Joint FAO/WHO Expert Committee on Food Additives. (1991).

40. Joint FAO/WHO Expert Committee on Food Additives. Evaluation of Certain Food Additives and Contaminants: Forty-Fourth Report of the Joint FAO/WHO Expert Committee on Food Additives. (1995).

41. Joint FAO/WHO Expert Committee on Food Additives. Evaluation of Certain Food Additives: Fifty-First Report of the Joint FAO/WHO Expert Committee on Food Additives. (1999).

